# Human fetal virtual histology with X-ray phase contrast imaging

**DOI:** 10.1101/2023.04.18.537316

**Authors:** Pierre-Louis Vérot, Laura Charnay, Clément Tavakoli, Shifali Singh, Laurène Quénot, Eva Solé Cruz, Monia Barnat, Sandrine Humbert, Marc Dommergues, Alexandra Durr, Christian Piolat, Pierre-Yves Rabattu, Emmanuel Brun

## Abstract

Human fetal analysis is of fundamental importance in understanding normal and pathological development. Congenital malformations and prenatal deaths are commonly linked to alterations of organs formation during embryonic and fetal life. Normal and pathological developments are not completely understood yet, because access to human fetal samples is difficult and exploratory means are scarce.

Here we show the first X-ray phase contrast images of *post mortem* human fetuses performed at the tissue scale. Imaging was performed on both an entire sample and isolated organs at higher resolution.

X-ray phase contrast imaging is based on the detection of refraction of a high energy X-ray beam after sample exposure. We were able to produce a tomographic imaging of human fetal tissues at the beginning of their second trimester at resolutions of 23 µm, 6 µm and 3 µm with high contrast on soft tissues.

Our results demonstrate how X-ray phase contrast imaging is a promising technique for human development analysis as it gives high-contrast and high-resolution results for soft and hard tissues.

We assume this technique to be a reference in the future of human development studies as it is conservative for rare and precious human specimens.

## Introduction

Congenital disorders are the cause of 25% of neonatal mortality in Europe and account for 295,000 neonatal deaths each year worldwide according to the World Health Organization[1]. Therefore, human development during its embryonic and fetal stages is a subject of high interest for medical applications. Most malformations requiring surgical or medical care during infancy find their origins during prenatal life[1]. Genetic and environmental factors can cause developmental defects inducing malformations or associations of malformations in a fetus[2]. Studying normal and pathological human fetuses is of uttermost importance in optimal care for survivors of the fetal period. A reproducible imaging, non-destructive that does not alter the sample would be of high interest. High-resolution and high-contrast 3 Dimensions (3D) imaging for human fetus development analysis is needed.

Conventional detailed histopathological study of fetal tissues requires tedious work in sectioning and preparation for microscope analysis[3]. Human prenatal specimens are rare and precious for developmental and anatomical studies.

A few years ago, an interactive 3D digital atlas[4] of human development was elaborated from the alignment of embryo sections during the first two months of development. This valuable work is well appropriate to perform an atlas but is too time-consuming and would require too much effort to extend to many cases for studying pathologies. More recently, high-resolution 3D images of the developing peripheral nervous, muscular, vascular, cardiopulmonary, and urogenital systems were obtained on human embryos and fetuses ranging from 6 to 14 weeks of gestation (WG)[5] using light-sheet microscopy combined with immunostaining. A combination of antibodies applied to each specimen allowed to perform a precise analysis of the developing systems at a cellular level. However, tissue shrinkage was observed on embryonic and fetal samples after sample preparation by solvents, limiting morphometric assessments. Moreover, the number of antibody combinations used on a single sample is limited[5]. Consequently, it would require a large number of samples to study a whole range of pathologies due to the technique’s destructiveness. To overcome this issue, Magnetic Resonance (MR) microscopy has been used for embryo imaging[6] allowing for 3D reconstruction. Nevertheless, this technique has limited contrast for hard tissues and has a limited resolution to evaluate the structures at the tissue scale. Nondestructive imaging technique allowing multiple resolutions from entire fetal bodies down to the functional unit of organs would be beneficial to understand the fetal development as a whole and the combination of congenital malformations leading to spontaneous abortions.

Recently X-ray Phase Contrast Imaging (PCI) has been introduced as a novel technique allowing a good contrast for different tissues[7] ranging from soft tissue such as breast tissue[8] or brain[9] to harder tissue such as cartilage and bone[10] without contrast agent. Contrary to conventional computed tomography based on the sole X-ray absorption, PCI is based on the detection of X-ray refraction by the tissues. The refraction index being up to 3 orders of magnitude greater than its counterpart, the absorption index, this modality provides a great contrast without the use of any contrast agent. Moreover, this technique has the great advantage of being applicable to a great range of resolutions from a few tens of nanometers[11] to patient radiographic imaging[12]. In detail, this technique uses a wave formalism of light propagation. It is now mainly available with synchrotron sources where, due to the coherence of the source, refraction through the objects induces interference patterns[13] on the image that are analyzed to retrieve the phase shift due to the sample. Some techniques allow to transfer the modality to conventional sources[14], [15] but their performances and resolution are limited.

To the best of our knowledge, only Kanahashi *et al*.[16], [17] imaged human embryos using this technique at a moderate spatial resolution and Garcia-Canadilla *et al*.[18] only on isolated fetal hearts.

Here we show the first results of *post mortem* human fetus imaging using Phase Contrast computed tomography at the tissular scale for a wide variety of organs together with an entire human fetus with the aim to demonstrate the potential of this conservative technique.

## Results

### Fetal full-body X-ray phase contrast imaging

Figure 1 shows multiple cross-sections of the entire fetus (13WG+3days) in axial coronal and sagittal sections of the thoracic, abdominal, and pelvic levels at the pixel size of 23 µm. Additionally, we present a 3D reconstruction of the bones as well as 3D segmentation of the heart pulmonary complex.

**Fig. 1:**
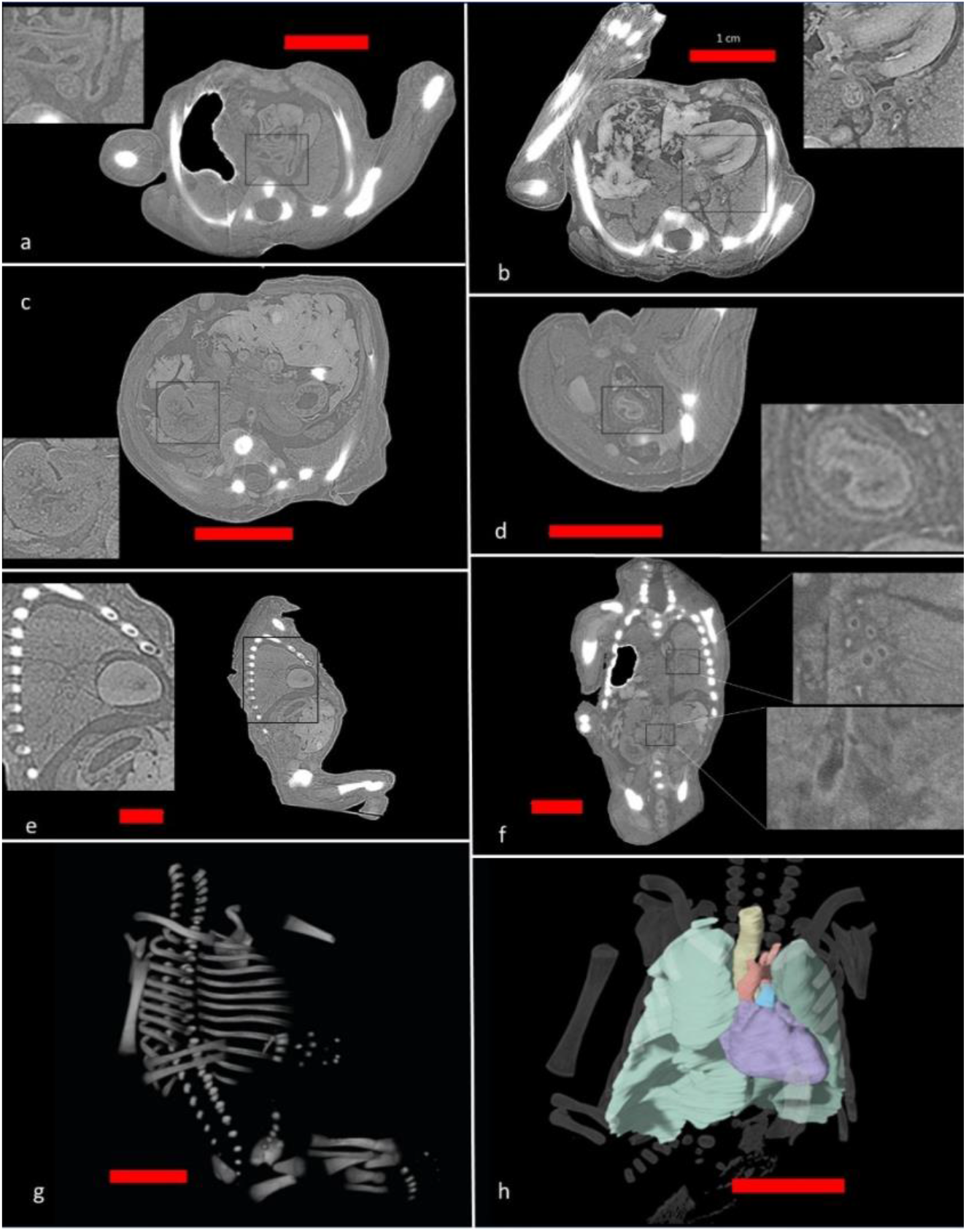
Tomographic phase contrast imaging of a 13WG+3days fetus and 3D rendering. **a** Transverse thoracic section with zoom on carina and ductus arteriosus **b** Transverse thoracic section with magnification on the left cardiac ventricle, thoracic esophagus, and left pulmonary hilum **c** Transverse abdominal section with enlargement on the right kidney **d** Pelvic cross-section with enlargement on the rectum **e** Fetal sagittal section with enlargement on the cardiac apex and the left lung with its oblique fissure **f** Coronal fetal section with enlargements on the left pulmonary hilum and the birth of the fetal renal arteries **g** 3D reconstruction of the fetal skeleton **h** 3D reconstruction of part of the intrathoracic contents: trachea in yellow, lungs in green, heart in purple, aortic arch and its branches in red, pulmonary artery trunk in blue with bone projections of a fetus. The red scale bar represents 1 cm.

The two-dimensional sections on a 13 gestational-weeks fetus allow us to visualize the fetal anatomy in detail. Figure 1a shows the carina in front of the thoracic esophagus. A large right pneumothorax caused by the medical intervention is visible and creates artifacts around its edge due to the large refraction at the interface air/tissues. Nevertheless, the ductus arteriosus connecting the aortic arch to the pulmonary trunk is visible. Figure 1b shows the fetal heart, the esophagus in the enlarged area with the mucosa in the shape of a Maltese cross on the thoracic region, and the lung parenchyma in its pseudo-glandular stage. The liver of this specimen is too damaged, and interpretation of its nature is impossible. Figure 1c presents the abdominal region. The magnification inset on the right renal region shows the renal pelvis, but it remains difficult at this resolution to individualize the renal structures (pedicle, glomerulus, etc.). The following figures present higher resolution images where nephrons are visible. Figure 1d shows the pelvis region where the different layers of the rectum can be visualized with a hyper signal on the mucosa. Figure 1e is a sagittal cross-section on the cardiac apex and the lung region. The oblique fissure of the left lung is visible as well as a part of the stomach body. The coronal section shown in Fig. 1f allows observing elements of the left pulmonary hilum and the origin of the renal arteries in the magnifications inset. Finally, 3D reconstructions of the skeleton (Fig. 1g) and segmented organs in the thorax are displayed (Fig. 1h).

### Isolated fetal organs visualization between 12 and 14 weeks of gestation

To evaluate the potential of PCI at the tissue scale, 6 µm and 3 µm scans of isolated organs were performed, and results are presented in the next sections and figures.

### Heart and lung

Figure 2 presents imaging results on the cardiopulmonary complex. The heart valves, myocardium, myocardial chambers, and the coronary arteries are well individualized, as shown in figs. 2a and 2b on a 12WG+2days fetus. 3D reconstruction after semi-automatic segmentation of the fetal coronary arteries is shown in the supplementary figures.

**Fig. 2:**
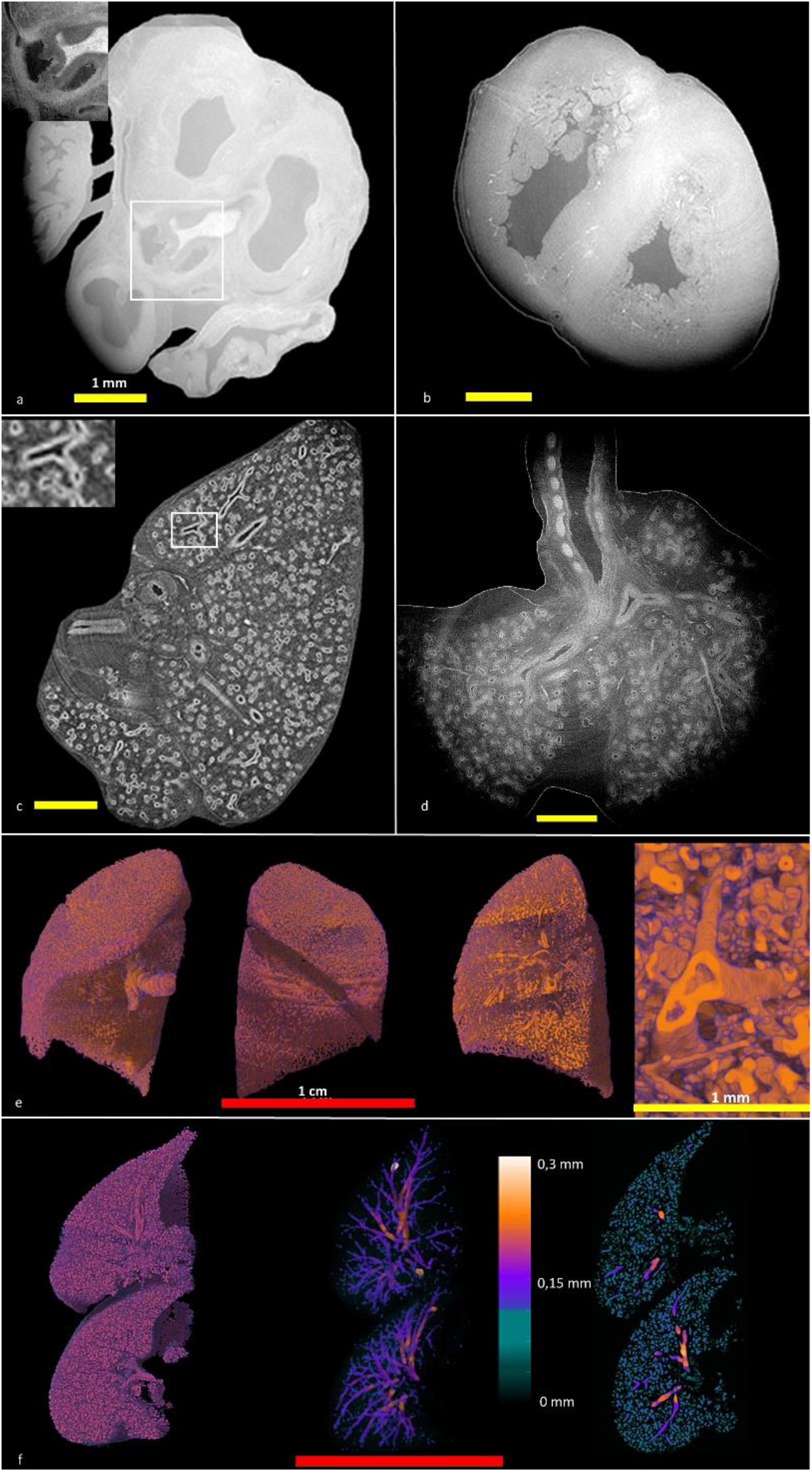
Tomographic phase contrast imaging of the heart and lungs of a fetus and 3D rendering of airways. **a**, Transverse section of fetal heart with enlargement on the aortic valve and sinus of Valsalva, birth of the coronary arteries. **b**, Cross section of fetal heart showing myocardium and ventricles. **c**, Cross-section of fetal lung at pseudoglandular stage with magnification on a sub-segmental bronchial division. **d**, Coronal section of fetal lungs at pseudoglandular stage showing trachea and main bronchi. **e**, 3D reconstruction of the bronchial and vascular tree of a right lung from a 12WG+2 days fetus with magnification on a bronchial division on the right. **f, Left:** 3D reconstruction of the bronchial and vascular tree, **Center:** 3D reconstruction with in false color structures ranging from 0.13 mm to 0.3 mm in diameter, **Right:** sagittal section in the volume. False color indicates a thickness ranging from 0.04 mm to 0.3 mm. The red scale bar represents 1 cm. The yellow scale bar represents 1 mm.

Figure 2c shows an axial section of a 13WG+6 days fetal lung. Figure 2d presents a coronal section of a 12WG+2 days lung with the trachea. Therefore, these lungs are in the pseudoglandular stage of development with bronchi multiplying in the parenchyma. The achieved contrast allows visualizing bronchi and blood vessels. Unfortunately, the contrast between the two structures is too limited to automatically segment separately the two trees. Nevertheless, we present in figs. 2e and 2f 3D rendering of these structures. Figure 2e presents the 3D rendering of the 12WG+2 days fetal lung in 3 different angles. The improved resolution owes to better distinguish the main bronchus in the left image and the inter lobar fissure. Finally, we present in the right part of figure 2e the same lung intersected by a plane where the subdivisions can be seen. Figure 2f presents a 3D representation of the two lungs of a 13WG+2days fetus. False color represents bronchi diameter measured by granulometry.

### Kidneys

The isolated kidneys used in this study were obtained from fetuses between 12 and 14 weeks of gestation. The maximum diameter of kidneys was 5.7 mm at 12 WG+2 days and 8.5 mm at 14 WG+ 2 days, with a volume increasing from 38.9 mm^3^ to 104 mm^3^ between these two weeks of development. Their appearance is lobulated, with irregular contours. Synchrotron phase contrast imaging allows visualization of the glomeruli, adjacent tubules, collecting tubes draining into the renal calyces, renal pelvis, and ureter (Fig. 3). Without any specific preparation or any injection of contrast medium, we can distinguish the urothelium in calix and in the ureter bordering the urinary excretory tract and glomeruli and tubules on the enlargement the kidney cortex (Fig. 3a). The arterial and venous blood vessels can be individualized (Fig. 3b) as well as their intraparenchymal branches. A 13WG + 2 days old fetal kidney was acquired at a resolution of 3 µm (Fig. 3c). This resolution allows the renal parenchyma to be observed in greater detail allowing individualization of glomeruli as shown on the enlargements. The improvement of resolution between 23 µm and 6 µm allows a better definition of the functional unit of kidneys. The improvement from 6 µm to 3 µm helps to visualize better, but the data size increase (up to 100 GB for one kidney) does not justify imaging all the samples at this resolution.

**Fig. 3:**
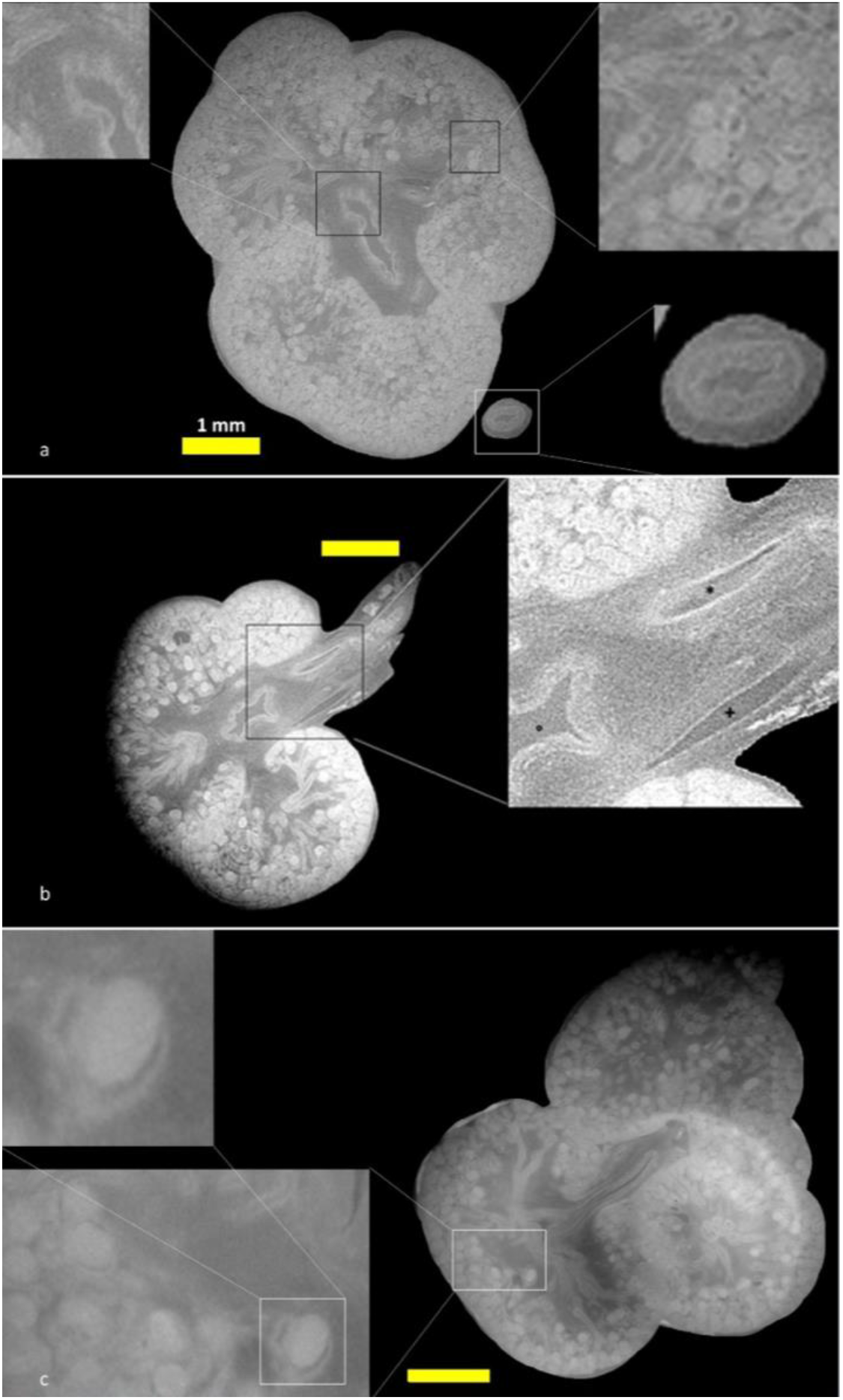
Tomographic phase contrast imaging of a 13 WG fetal kidney. **a**, Cross**-**section of a fetal kidney with enlargement on a calyx, the renal cortex, and the ureter. **b**, Cross-section of a fetal kidney with magnification on the hilum showing the renal artery *, vein +, and calyx °. **c**, 3 µm resolution cross-section of a fetal kidney with magnification on the parenchyma and an individualized glomerulus. The yellow scale bar represents 1 mm.

### Eyes

The results shown in Fig. 4 are from a fetal eye of 13WG. The eye is 5 mm in diameter. The cornea, uvea, lens, sclera, retina, and optic nerve are clearly identifiable. The hyaloid canal located in the vitreous body is hyperdense in phase contrast; it contains the hyaloid artery permeable during fetal life. The nerve fibers are visible in the enlarged area.

**Fig. 4:**
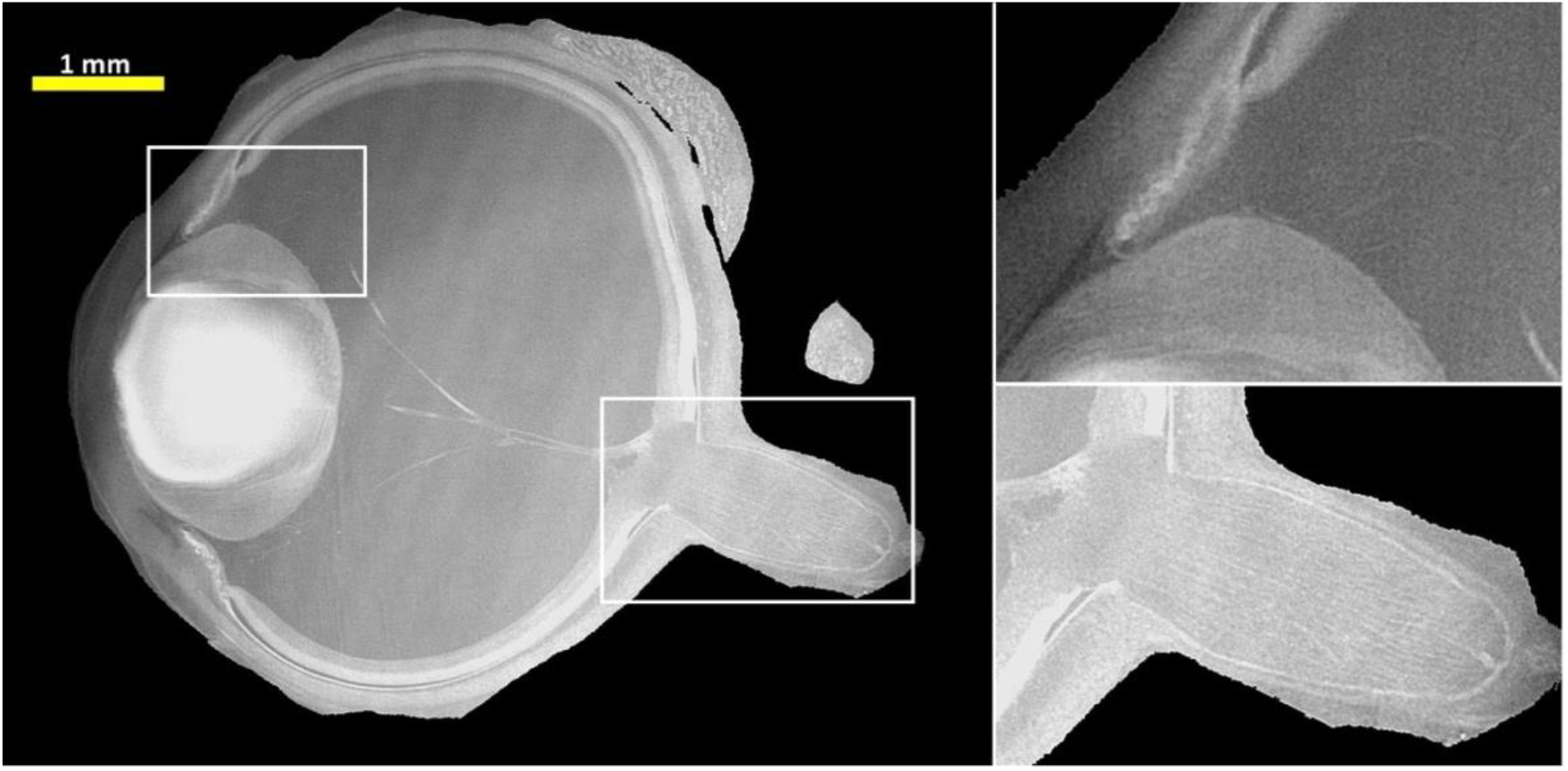
Tomographic phase contrast imaging of a 13 week of gestation fetal eye with magnification on the ciliary body and the optic nerve. The yellow scale bar represents 1 mm.

### Esophagus and intestines

The digestive tract is illustrated in Fig. 5, with the laryngopharynx in Fig. 5a composed of the thyroid cartilage in front and the arytenoid cartilages in the center where the vocal folds - the origin of the vocal cords - are docked. The fibers of the pharyngeal muscles are visible, especially the thyroarytenoid muscle in Fig. 5a. The esophagus is presented in Fig. 5b and its mucosa with a high density relative to the muscular in phase contrast imaging, shaped like a Maltese cross. The esophagus has a diameter of 1 mm. The small intestine of the same 13WG + 1-day-old fetus is shown in Fig. 5c. PCI allows the visualization of the intestinal mucosa, as well as the mesentery containing the mesenteric vessels. The intestinal mucosa is denser than the muscular and serosa and already forms the villi, which gives a large contact surface with the intestinal lumen. The entanglement of the jejunal and ileal intestinal loops can be seen in the 3D volume projection in Fig. 5d. The mesentery curtain is visible and magnified, allowing the relief of the mesenteric vessels within it to be discerned.

**Fig. 5:**
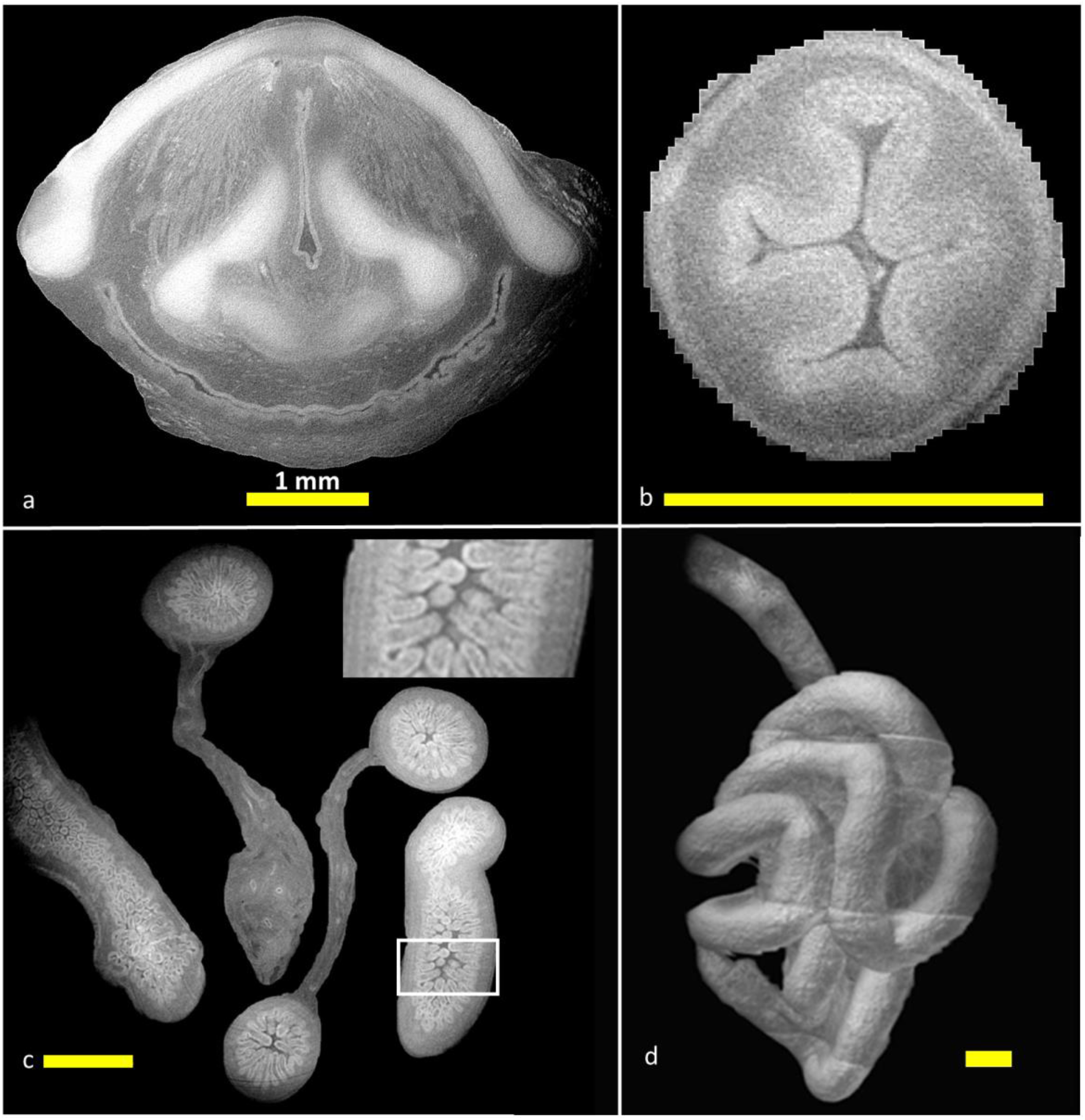
Tomographic phase contrast imaging of the fetal digestive track at 13 weeks of gestation. **a** Cross-section of a fetal laryngopharynx. **b**, Cross-section of a fetal thoracic esophagus. **c**, Cross-section of a fetal intestine with magnification on intestinal villi. **d**, 3D reconstruction of the fetal gut showing the mesentery. The yellow scale bar represents 1 mm.

## Discussion

We performed, for the first time, PCI-CT of both an entire fetus and isolated human organs without deforming or degrading them. The entire 13 weeks of gestation fetus was imaged at a pixel size of 23 µm, and the isolated organs were imaged at both 6 and 3µm pixel size. We present results of this imaging technique on diverse organs ranging from soft tissue kidneys to hard tissue such as cartilage or bone at a tissue level without any degradation on the sample. So far, nondestructive studies found in the literature were investigating fetal anatomy with Magnetic Resonance Imaging (MRI)[19] or Phase Contrast Imaging (PCI)[16] at a lower resolution. The resolution and the contrast achieved in this work allowed us to study an entire fetus on all the different tissue types at the tissue level, allowing one to study the functional unit of fetal organs and their morphological evolution without the use of contrast agent. Indeed compared to diffuse iodine-based CT[20] that induce a certain alteration of tissue, our method is completely non invasive and conventional foetopathology could be performed after imaging.

We were able to highlight details in the structure of kidney parenchyma with sufficient resolution to individualize the components of the nephrons. The focus of this article was to explore all different organs and their contrast. In principle, a high-quality morphological analysis similar to the one published by Zdora *et al*.[21] could also have been made on a human fetal kidney. Thanks to the capability of imaging a whole fetus, one could describe kidney migration during fetal development with multiple samples or the partition of the esophagus from the trachea, or the segmentation of the lungs, all of which have fetal pathological implications.

Studying the development of human fetal lung is a current challenge[22]. The development of the respiratory tree is divided into 5 consecutive phases: embryonic, pseudoglandular, ductal, saccular, and alveolar. In this study we could only acquire lungs at the pseudoglandular stage due to the samples we had. Our images appear to be of sufficient quality to follow lung development in the pseudoglandular stage as we could study in 3D the bronchial tree using artificial intelligence for the automatic segmentation of isolated components of interest (bronchi, vessels, etc.). This technique could make possible the description of the pulmonary circulation evolution and, in particular, the transition between the systemic and pulmonary vascularization.

Cardiac defects account for approximately 20% of congenital malformations in live births[1]. Fetal heart tissue has recently been studied using phase contrast imaging along with post-processing treatments allowing precise analysis of myocardial micro- and macro-structures[18], in a 20-weeks of gestation fetus with a heart defect, compared to a healthy 19-weeks of gestation fetal heart. Here we were able to analyze the heart tissue of 11 to 13-weeks of gestation fetuses in this work. The coronary vessels were well individualized as well as the myocardium and valvular structures. In the complete fetus, the ductus arteriosus was individualizable among the large vessels.

The mesenteric vascularization of the intestine could be highlighted in this work, its internal structure could also be visualized. We were able to obtain images of the intestinal mucosa with its villi, already present during the second trimester. Applied to a complete embryo, the PCI technique could be used to study this implantation, but the whole sample used in this study was not of sufficient quality. A study of the reintegration of the primitive bowel loop by PCI was performed by Kanahashi et al.[17], showing that the reintegration of the loop is independent of the fetal liver size. The imaging study of the length and growth of the fetal bowel in the second trimester is a current issue[23], [24]. This issue is known to have direct essential clinical application for the consequences of intestinal damage in premature children. Probably the study of mesenteric vascularization with PCI could allow studying the pathophysiology of intestinal atresia or intestinal malposition[25].

Some limitations of this work deserve to be mentioned, first the fetal samples used were partly degraded due to the medical intervention. Then, we could not study the evolution of fetal development over time and the number of samples was insufficient to allow comparative quantitative studies. Nevertheless, we were able to obtain sufficient human fetal material to explore many developing organs and systems.

To conclude, this study demonstrates the feasibility of fetal phase contrast imaging and presents the range of potential applications of this technique in the study of normal and pathological human development. This imaging technology could be suitable for diagnosis of fetal pathologies instead of the usual fetopsy when High resolution phase contrast imaging will no longer depend on high energy synchrotron radiation source[26].

## Methods

Partial and complete human fetal samples used in this work had been previously used for a neurodevelopmental study on Huntington’s disease[27], which essentially focused on the brain. The research use of these samples was authorized by the French Biomedicine Agency (authorization n°PFS17-001; 24/01/2017) accordingly to the Helsinki declaration. We were able to obtain a nearly complete fetus with a gestational age of 13 WG + 3 days, and 9 other fetuses with only certain portions or isolated organs. These samples were placed in hermetically sealed tubes filled with PBS (phosphate-buffered saline) solution without further treatment before the acquisitions.

**Table 1.** Characteristics of 10 analyzed fetal samples. ALS= Amyotrophic Lateral Sclerosis; HD= Huntington Disease

| Gestational age<br>(WG+days) | Associated pathology | Organs |
| --- | --- | --- |
| 14+1 | ALS | Kidneys, adrenal gland, trachea, heart |
| 12+3 | None | Lower limb, hands |
| 13+2 | None | Lung, kidney |
| 13+2 | None | Eyes, feet, kidneys, heart |
| 12+2 | HD | Lower limb, fore-arm, hand, kidneys, adrenals, heart, trachea, lungs and eye |
| 13+2 | HD | Kidneys, foot, wrist, hand, eyes |
| 11+6 | None | Eye, lung, fore-arm, hand |
| 13+3 | None | Entire fetus |
| 13+6 | HD | Lung, kidney, intestine, adrenal gland |
| 13+1 | HD | Kidney, intestine, hand, oesophagus and trachea |

### X-ray Imaging Experiment

We used a so-called Propagation-based Phase Contrast Imaging set-up. The experiment was performed at the biomedical beamline (id17) of the ESRF, the European Synchrotron. The energy was set to 33 keV by using a Laue Laue monochromator. The sample to detector distance was 4.5 m. The X-ray detectors used were a sCMOS detector coupled with different optics resulting in a pixel size of 3, 6, and 23 µm. We collected 3600 projections around 360 degrees. The total acquisition time for one rotation was 3 min. Due to the limited height of the synchrotron beam, the samples were vertically translated in order image them entirely. The maximal acquisition time for the biggest sample at the maximal resolution was 2.5 hours. A single distance phase retrieval algorithm [13] was used with a fixed refraction/absorption ratio of 1200. CT reconstruction was performed using PyHST software [28].

### Data Analysis

Once reconstructed, the images were post-processed (stitching, ring artefact reduction) with a home-made software [29]. The 3D visualizations were made with dragonfly (Object Research Systems, Montreal, QC, Canada). The machine learning method used was implemented in the Explorer software (Reactiv’IP, Grenoble France); it consists of a random forest with different image filters. Algorithms in python and Artificial Intelligence Models can be provided upon reasonable request.

## Acknowledgement

We are deeply grateful to the parents for giving their full consent and allowing us to do this study. We acknowledge the ESRF for granting beamtime (MD1217).

## Author contributions statements

PLV, ESC, MB, SH, MD, AD, CP, PYR, and EB conceived the work and design the study. PLV, LC CT SS, LQ and EB performed the acquisition and analysis. The data were interpreted by everyone. CT, SS LQ and EB created the software popcorn for the data analysis. The manuscript has been drafted and revised by all the authors.

